# Multimodal brain-age prediction and cardiovascular risk: The Whitehall II MRI sub-study

**DOI:** 10.1101/2020.01.28.923094

**Authors:** Ann-Marie G. de Lange, Melis Anatürk, Tobias Kaufmann, James H. Cole, Ludovica Griffanti, Enikő Zsoldos, Daria Jensen, Sana Suri, Nicola Filippini, Archana Singh-Manoux, Mika Kivimäki, Lars T. Westlye, Klaus P. Ebmeier

## Abstract

Brain age is becoming a widely applied imaging-based biomarker of neural aging and potential proxy for brain integrity and health. We estimated multimodal and modality-specific brain age in the Whitehall II MRI cohort using machine learning and imaging-derived measures of gray matter morphology, diffusion-based white matter microstructure, and resting state functional connectivity. Ten-fold cross validation yielded multimodal and modality-specific brain age estimates for each participant, and additional predictions based on a separate training sample was included for comparison. The results showed equivalent age prediction accuracy between the multimodal model and the gray and white matter models (R^2^ of 0.34, 0.31, and 0.31, respectively), while the functional connectivity model showed a lower prediction accuracy (R^2^ of 0.01). Cardiovascular risk factors, including high blood pressure, alcohol intake, and stroke risk score, were each associated with more apparent brain aging, with consistent associations across modalities.

## 1. Introduction

In older age, the human brain undergoes structural changes including reductions in brain volume, cortical thinning, and decline in white-matter microstructure [1], and large-scale resting-state networks become less segregated [2, 3]. Aging-related changes in brain structure and functional connectivity are associated with decreased cognitive performance in domains including memory and processing speed [2, 4, 5], and comprise an increased risk for neurodegenerative disorders such as dementia [6]. Although the senescent deterioration of the brain is well-known, older populations are characterised by substantial variation in neurobiological aging trajectories [1], and recent neuroimaging studies have focused on developing potential markers for brain aging [7, 8]. Brain-age prediction based on machine-learning algorithms estimates an individual’s ‘brain age’ using structural and functional brain characteristics derived from magnetic resonance imaging (MRI) [9, 10, 11, 8]. Subtracting chronological age from estimated brain age provides an estimate of brain aging, the *brain-age delta*. For instance, if a 70 year old individual exhibits a brain-age delta of +5 years, their typical aging pattern resembles the brain structure of a 75 year old, i.e. their estimated brain age is older than what is expected for their chronological age [11]. Individual variation in delta estimations are associated with a range of cognitive and biological measures [10, 11, 12, 13, 14, 15, 16, 17], including cardiovascular health [14], and differences in brain age have been established between patient groups and healthy controls: individuals with conditions such as Alzheimer’s disease, multiple sclerosis, epilepsy, and psychiatric disorders show on average older brain age relative to their chronological age [9, 18, 19, 20, 21, 22]. Longitudinal studies have documented highly reliable brain age prediction in stroke patients [23], and accelerated brain aging in patients with schizophrenia and multiple sclerosis [21, 24, 25]. Combined with studies on the association between brain-age delta and biomedical factors in healthy population cohorts [13, 15], the documented reliability and clinical sensitivity supports the utility of brain-age estimation as a candidate biomarker for neurological senescence and disease [8].

Modality-specific brain age models (based on e.g. gray and white matter separately) provide information about tissue-specific aging processes [26, 27]. For instance, imaging-derived measures of gray matter are known to detect cortical atrophy in older age-groups [28], while changes in diffusion MRI measures reflect age-related decline in white matter microstructure, as well as white matter lesions, which are more prevalent in aging, relative to young, populations [29]. Functional MRI (fMRI) measures are indicative of brain network connectivity, which may change with advancing age [2, 3]. Cardiovascular risk factors may influence these neural aging processes differently [26, 27, 30], and in a recent Whitehall II (WHII) MRI study using voxelwise analyses, allostatic load, metabolic syndrome, and multifactorial stroke risk predicted gray matter density measured decades later, while only cumulative stroke risk measured by the Framingham stroke risk score [31] predicted white matter integrity in terms of fractional anisotropy and mean diffusivity [32].

In this study of the WHII MRI cohort (N = 671), we investigated whether machine learning using neuroimaging data could produce reliable biomarkers of brain aging, and whether cardiovascular risk factors including blood pressure and alcohol intake, and cumulative risk as indicated by the Framingham stroke risk score were associated with modality-specific brain-age markers. We estimated brain age using ten-fold cross validation in separate models based on I) gray matter (GM) measures, II) white matter (WM) measures derived from diffusion tensor imaging (DTI) and white matter hyperintensites (WMH), III) functional connectivity measures derived from resting state fMRI (rs-fMRI), and IV) a multimodal model that included all of the brain measures. A detailed description of the methodology is provided below.

## 2. Materials and Methods

### 2.1. Sample

The WHII study was established in London in 1985, and included an initial cohort of 10,308 civil servants. Between 2012 and 2013, 6035 individuals participated in the Phase 11 assessment, from which a random sample of 800 participants was enrolled in an MRI sub-study including brain scans and biomedical assessments [33] (www.psych.ox.ac.uk/research/neurobiology-of-ageing/research-projects-1/whitehall-oxford). The current sample was drawn from the WHII MRI sub-study, and included 715 participants with multimodal MRI data. Forty-four participants were excluded based on neurological disease and incidental MRI findings, yielding a final sample of 671 participants. Sample demographics are provided in Table 1.

**Table 1:** Sample demographics. Age range (mean age ±standard deviation = 54.70 ±7.29), percentage male (M) and female (F) participants, percentage with white (W) and non-white (NW) ethnic background, and percentage with educational qualifications U = university degree, PG = post-graduate / masters / PhD, Pr = Professional qualifications, A = A levels or equivalent, O = O levels or equivalent, N = No qualifications.

| N | Age range | Sex % | Ethnicity % | Educational qualification % |
| --- | --- | --- | --- | --- |
| 671 | 60.43 - 84.58 | M79 \| F21 | W95 \| NW5 | U27 \| PG22 \| Pr12 \| A18 \| O14 \| C5 \| N3 |

### 2.2. MRI data acquisition and processing

MRI data were acquired using a 3 Tesla Siemens Magnetom Verio (n. of participants = 473) with a 32-channel receive head coil (between April 2012 – Dec 2014) and, following scanner updates, a 3 Tesla Siemens Magnetom Prisma (n. of participants = 198) with a 64-channel head-neck coil (June 2015 – Dec 2016). MRI images for all participants were processed using the analysis pipeline described in [33], including automated surface-based morphometry and subcortical segmentation as implemented in FreeSurfer 6.0 [34], and residualized with respect to scanner, relative head motion during the acquisition of rs-fMRI images [35, 36], intracranial volume (ICV [37]), sex, and ethnic background using linear models.

#### 2.2.1. Gray matter

In line with recent large-scale implementations [16, 19], we utilized a fine-grained cortical parcellation scheme [38] to extract cortical thickness, area, and volume for 180 regions of interest per hemisphere, in addition to the classic set of subcortical and cortical summary statistics from FreeSurfer [34]. This yielded a total set of 1118 structural brain imaging features (360/360/360/38 for cortical thickness/area/volume, as well as cerebellar/subcortical and cortical summary statistics, respectively). To exclude potential outliers, we tested for values that were ± 4 standard deviations (SD) away from the average on the global MRI measures mean cortical or subcortical gray matter volume. No outliers were identified based on the general T1-derived measures.

#### 2.2.2. White matter microstructure

Global and tract-specific estimates of fractional anisotropy (FA), mean diffusivity (MD), axial diffusivity (AD), radial diffusivity (RD) and mode of anisotropy (MO) were calculated for each individual. In accordance with established methods [39], tract-specific estimates of each DTI metric were derived using 48 standard-space masks available from the ICBM-DTI-81 White-Matter Labels Atlas [40, 41], producing a total of 245 DTI features. Global WMH volumes were automatically extracted from FLAIR images with Brain Intensity AbNormality Classification Algorithm (BIANCA) [42]. To avoid scanner-specific biases in these estimates, BIANCA was initially trained with WMH masks manually delineated in a sub-sample of individuals scanned on the Prisma (n = 24) and Verio (n = 24) scanners and an independent sample from the UK Biobank study (n = 12) [43]. Subjects with values ± 4 SD away from mean FA or mean MD were excluded, yielding a total of 668 included subjects.

#### 2.2.3. Functional connectivity

Spatial maps of large-scale resting state networks were derived by applying MELODIC group Independent Component Analysis (group-ICA, n of components = 25) [44] to the rs-fMRI images of 678 individuals of the Whitehall II MRI sub-study. All non-artefactual group-ICA components (n = 19) and subject-specific rs-fMRI timeseries (extracted with dual regression [45, 46]) were used in FSLNets (https://fsl.fmrib.ox.ac.uk/fsl/fslwiki/FSLNets) in order to generate a 19×19 matrix representing subject-specific correlations between each pair of networks. Partial correlations (derived using L2 Regularization, setting rho = 0.01 in Ridge Regression option in FSLNETS) were examined in the present study, as these estimates allow for a more direct estimate of the connectivity between each pair of nodes [47]. These partial correlations were then z-transformed using Fisher’s transformation. Subsequently, the upper triangle of the matrix was converted into a row vector, producing a total of 171 rs-fMRI features for each participant. Subjects with values ± 4 SD away from the average on any of the fMRI measures were excluded, yielding a total of 651 included subjects.

### 2.3. Brain age prediction

#### 2.3.1. K-fold cross-validation

The XGBoost regressor model https://xgboost.readthedocs.io/en/latest/python) was used to run the brain-age prediction, including an algorithm that has been used in recent large-scale brain age studies [16, 19]. Parameters were set to maximum depth = 3, number of estimators =100, and learning rate = 0.1 (defaults). Age was estimated in ten-fold cross validations yielding a multimodal brain-age estimate for each individual, as well as modality-specific brain-age estimates based on gray matter, white matter, and functional connectivity measures. To investigate the prediction accuracy, correlation analyses were run for predicted versus chronological age, and R^2^, root mean square error (RMSE), and mean absolute error (MAE) were calculated for each model. After removing outliers for each respective modality, the multimodal model included 649 subjects. For this model, average RMSE was calculated from a cross validation with ten splits and ten repetitions, and compared to a null distribution calculated from 1000 permutations. The result is shown in Figure 1.

**Figure 1:**
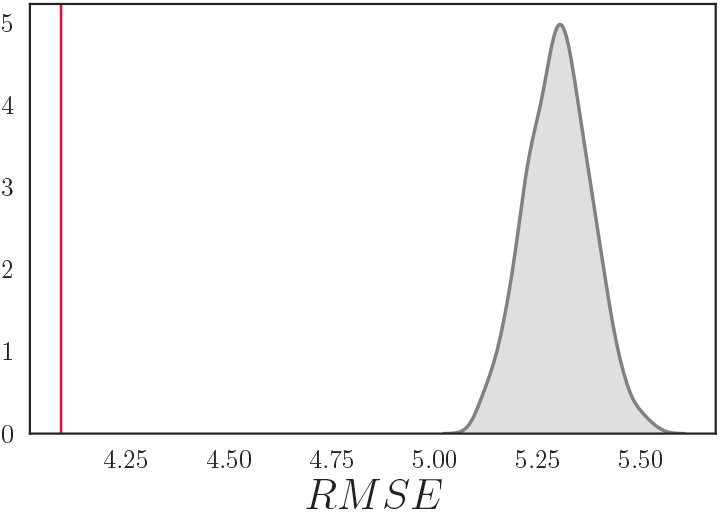
The mean ± SD root mean square error (RMSE) for the multimodal brain age model was 4.09 ± 0.32 based on the cross validation (red vertical line). The null distribution calculated from 1000 permutations is shown in grey, with a mean ± SD of 5.30 ± 0.07. The number of permuted results from the null distribution that exceeded the mean from the cross validation was 0 (*p* < 0.001).

#### 2.3.2. Data quality analyses

To investigate effects of including versus excluding low-quality imaging data in the models, we trained two separate models for each modality (with and without low-quality data) on 60% of the sample, and tested them on the remaining 40% of the data. The categorization of data was based on manual quality checks and images were classified as “poor quality” if they contained evidence of excessive motion and/or scanner-related artefacts. For diffusion weighted images, images with > 5 volumes missing from their scans (which were excluded due to containing 10 or more outlier slices) were also labelled as “poor quality”. The percentage of subjects with low-quality data in the training sets was 9% for the gray matter model, 12% for the white matter model, 13% for the RS functional connectivity model, and 11% for the multimodal model. To test for differences in model performance before and after excluding low-quality data, we used a *Z* test for correlated samples [48]:

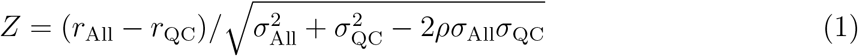

where “All” represents the full sample, “QC” represents the sample with low-quality data removed, the *r* terms represent the Pearson’s correlation coefficients of predicted versus chronological age, the *σ* terms represent their errors, and *ρ* represents the correlation between the two sets of model predictions.

#### 2.3.3. External training sets

As a cross check, we estimated gray-matter based brain age in the current sample using an external model trained on a 27,200 subjects from UK Biobank, with a mean age ±SD of 55.40 ±7.46, and 48% male / 52% female subjects (*External model 1*). The model contained the same 1118 structural MRI variables as those included in the WHII k-folding model (see section 2.2.1). The MRI data were residualized with respect to scanning site, data quality and motion using Euler numbers [49] extracted from FreeSurfer, ICV, sex, and ethnic background using linear models. Subjects with known brain disorders were excluded based on ICD10 diagnose (chapter V and VI, field F and G, excluding G5, http://biobank.ndph.ox.ac.uk/showcase/field.cgi?id=41270). 43 subjects with values ± 4 SD away from the average on the measures mean cortical or subcortical gray matter volume were excluded, yielding a total of 27,157 subjects. The correlation between the brain-age estimates based on this training set and the estimates based on the k-folding procedure for gray matter was *r* = 0.49, *p* < 0.001, 95% confidence interval (CI) = [0.43, 0.54]. When applied to the WHII dataset, the RSME was 10.28, and the correlation between predicted and chronological age was *r* = 0.49, *p* < 0.001, 95% CI = [0.43, 0.54].

To test the performance of a sex-specific external model with a larger age range, we applied an additional gray matter model trained on a separate sample of 35,474 individuals aged 3–89 (mean age ±SD = 47.30 ±17.10) [19] (*External model 2*). The model was derived from https://github.com/tobias-kaufmann/brainage. For this analysis, the WHII data were corrected for scanning site, estimated in-scanner head motion, ICV, and ethnic background. The external model was trained on men and women separately [19], and applied to each sex in the WHII sample. The correlation between the brain-age estimates based on this training set and the estimates based on the k-folding procedure for gray matter was *r* = 0.64, *p* < 0.001, 95%CI = [0.59, 0.68]. When applied to the WHII dataset, the RSME was 9.58, and the correlation between predicted and chronological age was *r* = 0.47, *p* < 0.001, 95% CI = [0.41, 0.53]. As the two external models showed similar performance, we included only the sex-specific *external model 2* with larger age range in the subsequent analyses, herein referred to as “the external model”.

### 2.4. Brain age delta and biomedical measures

To investigate associations with biomedical variables, the brain-age delta (predicted age – chronological age) was used as a measure of apparent brain aging. Clinical measures included systolic and diastolic blood pressure, alcohol intake measured by units per week (see [50] for details), and the Framingham stroke risk score [31], which includes cardio-metabolic measures, smoking habits, diabetes, sex, and age (see [32] for full description). Blood pressure and alcohol intake were corrected for sex, ethnic background, and educational level using linear models. Stroke risk score was corrected for ethnic background and educational level, as sex was already accounted for in the score. 624 subjects had data on all biomedical variables as well as demographic variables. Subjects with values ± 4 SD away from the average on any of the variables were excluded from the analyses, yielding 616 subjects in total. The mean ±SD for each measure is shown in Table 2.

**Table 2:** Mean ±standard deviation on each of the biomedical measures.

| N | Systolic BP | Diastolic BP | Alcohol intake | Framingham Stroke Risk score |
| --- | --- | --- | --- | --- |
| 616 | 140.52 ± 16.48 | 77.17 ± 10.47 | 14.55 ± 13.47 | 11.04 ± 5.46 |

To obtain a direct comparison of *β* values, both the brain-age deltas and the clinical variables were standardized (subtracting the mean and dividing by the SD) before a series of multiple regressions were run. In order to adjust for a frequently observed bias in brain-age prediction leading to an overestimation of brain age in younger subjects and an underestimation of brain age in older subjects [14, 15, 30, 51, 52], chronological age was included as a covariate. Correction for multiple comparisons was performed using false-discovery rate correction [53]. The statistical analyses were conducted using Python 3.7.0.

## 3. Results

### 3.1. Brain age prediction

Figure 2 shows the correlations between predicted age and chronological age for each of the models. The multimodal, gray matter, and white matter models showed consistent performance with explained variance (R^2^) ±standard error (SE) of 0.33 ±0.02, 0.31 ±0.02 and 0.31 ±0.03, respectively, while the functional connectivity model showed lower prediction accuracy with an R^2^ ± SE of 0.02 ±0.04. The explained variance of the predictions based on the external gray matter model was 0.22 ±0.03. The accuracy of each model’s prediction measured by R^2^, adjusted R^2^, RMSE, and MAE are shown in Table 3. As a follow-up analysis, we re-ran the multimodal model without the rs-fMRI data to test for prediction improvements. The results showed comparable performance with and without the rs-fMRI data (RMSE = 4.14 vs 4.17, *z* = −1.14, *p* = 0.26). Figure 3 shows the proportion of variance in chronological age explained by each of the models. Figure 4 shows the correlations between the multimodal and modality-specific brain-age deltas. The delta values were first corrected for age-bias [14, 15, 30, 51, 52] using linear models, and the residuals were used in the correlation analyses. Excluding low-quality data improved the multimodal model (*z* = 2.07, *p* = 0.04), but none of the other models, as shown in Table 4 and 5.

**Figure 2:**
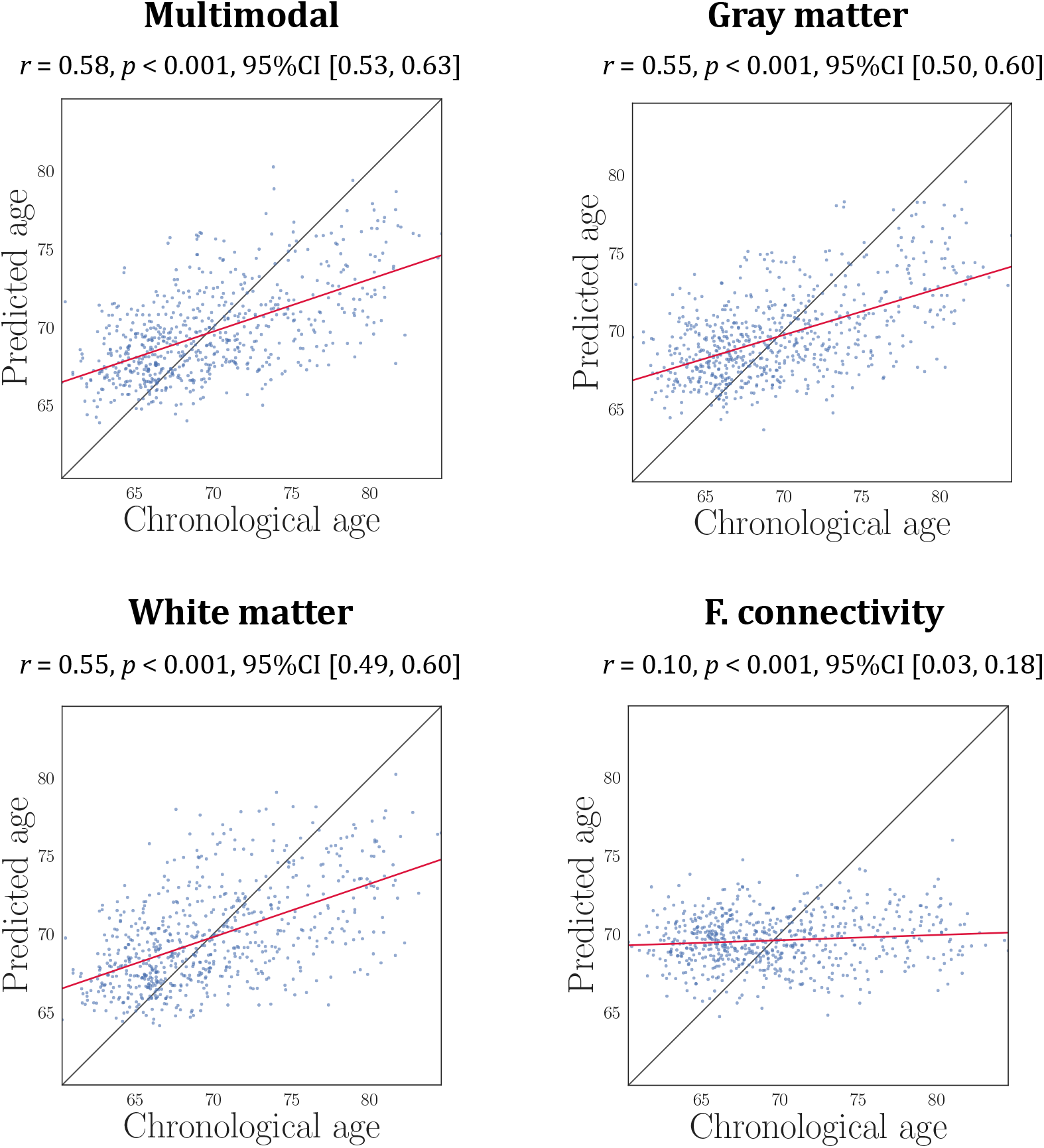
The association between predicted and chronological age for each of the models is shown by the red line. The black line illustrates a perfect relationship between x and y.

**Table 3:**
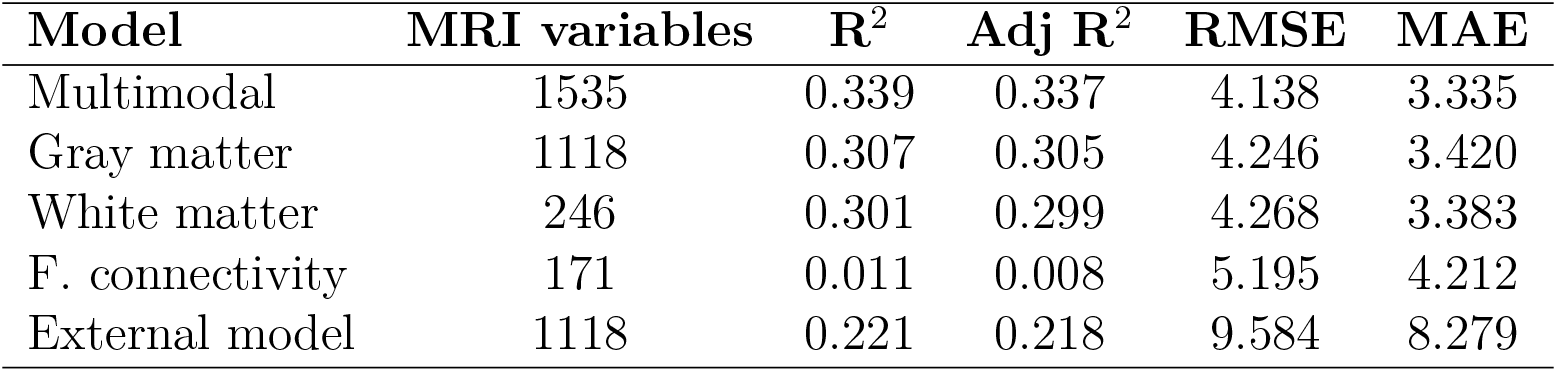
Number of MRI variables, R^2^, adjusted R^2^, root mean square error (RMSE), and mean absolute error (MAE) for each of the brain-age models. RMSE and MAE are reported in years.

**Figure 3:**
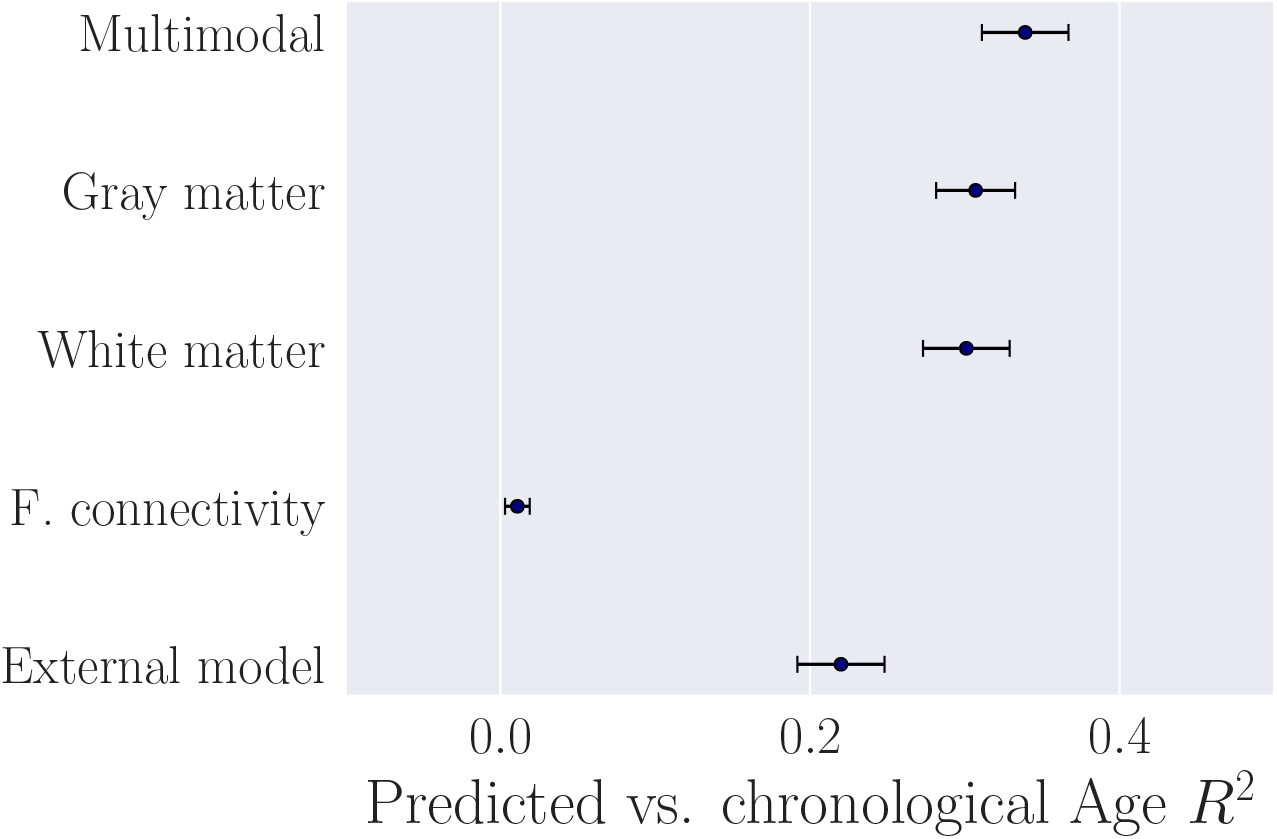
The proportion of variance in chronological age explained (R^2^±standard error) by the multimodal model, the modality-specific models, and the external model based on a separate training sample.

**Figure 4:**
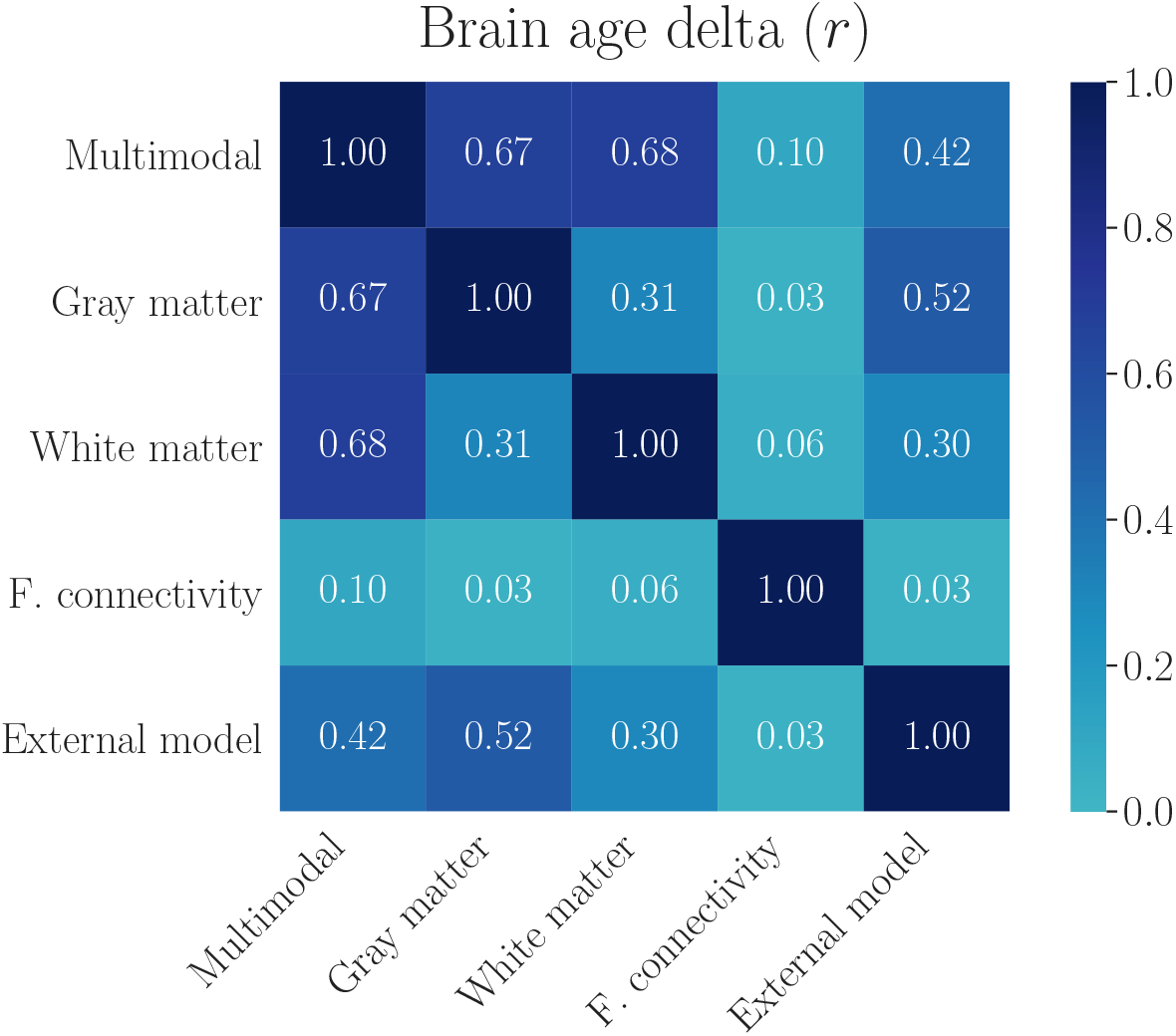
The correlations (Pearson’s *r*) between brain-age deltas of the multimodal model, the modality-specific models, and the external model based on a separate training sample. The delta values are corrected for chronological age.

**Table 4:** The correlation (Pearson’s *r*) between predicted and chronological age based on training sets with and without low-quality data applied to the same test set. 95% confidence intervals are indicated in square brackets. *Z* represents the difference in r values expressed in standard deviations, accounting for the correlated samples (Eq.1.)

| Model | All data included | QC data excluded | $Z$ | $p$ |
| --- | --- | --- | --- | --- |
| Multimodal | $r = 0.54, [0.44, 0.62]$ | $r = 0.57, [0.48, 0.64]$ | 2.07 | 0.04 |
| Gray matter | $r = 0.56, [0.47, 0.64]$ | $r = 0.59, [0.50, 0.66]$ | 1.70 | 0.09 |
| White matter | $r = 0.51, [0.42, 0.59]$ | $r = 0.50, [0.40, 0.58]$ | -0.72 | 0.47 |
| F. connectivity | $r = 0.08, [-0.04, 0.20]$ | $r = 0.07, [-0.05, 0.19]$ | -1.57 | 0.12 |

**Table 5:**
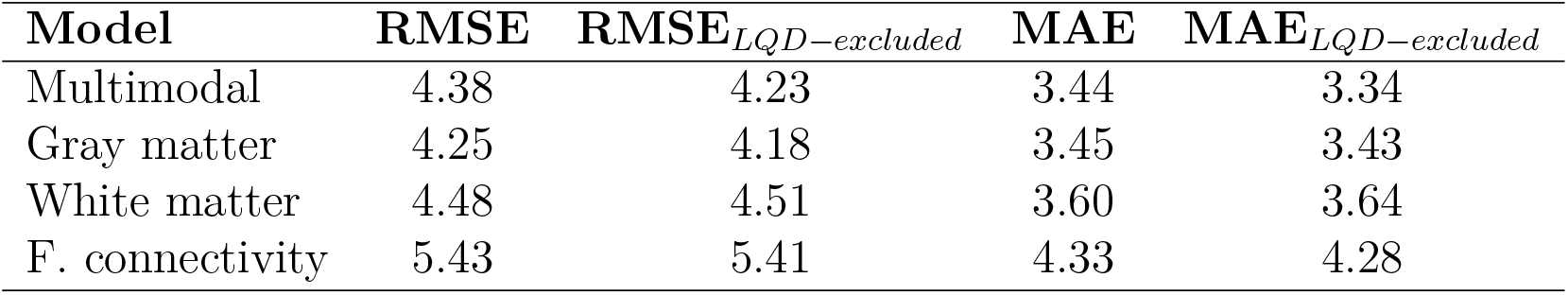
Root mean square error (RMSE) and mean square error (MAE) for the predictions based on training sets with and without low-quality data (LQD) applied to the same test set. RMSE and MAE are reported in years.

| Model | RMSE | RMSE<sub>LQD-excluded</sub> | MAE | MAE<sub>LQD-excluded</sub> |
| --- | --- | --- | --- | --- |
| Multimodal | 4.38 | 4.23 | 3.44 | 3.34 |
| Gray matter | 4.25 | 4.18 | 3.45 | 3.43 |
| White matter | 4.48 | 4.51 | 3.60 | 3.64 |
| F. connectivity | 5.43 | 5.41 | 4.33 | 4.28 |

### 3.2. Biomedical predictors

The associations between blood pressure (BP), alcohol intake, and stroke risk and brain-age deltas are shown in Figure 5 and Table 6. The associations were consistent across models. To test for non-linear relationships, polyfits including both a linear and a quadratic term (γ) were run for the multimodal brain age and each variable. No non-linear associations were found (Systolic BP: γ = 0.002±0.028, *p* = 0.948; Diastolic BP: γ =—0.003±0.028, *p* = 0.928;; Alcohol intake: γ = 0.023±0.028, *p* = 0.413; Framingham stroke risk score: γ = 0.004±0.028, *p* = 0.887).

**Table 6:** Relationships between biomedical variables and brain-age delta for each modality, including age as a covariate. *p*-values are reported before and after FDR correction. Corrected *p*-values below 0.05 are marked with an asterisk.

| Model | $\beta$ | SE | $t$ | $p$ | $p_{corr}$ |
| --- | --- | --- | --- | --- | --- |
| **BP Systolic** |  |  |  |  |  |
| Multimodal | 0.04 | 0.02 | 1.79 | 0.073 | 0.112 |
| Gray matter | 0.00 | 0.02 | 0.22 | 0.826 | 0.826 |
| White matter | 0.05 | 0.02 | 2.00 | 0.046 | 0.077 |
| F. connectivity | 0.03 | 0.01 | 2.78 | 0.006 | 0.022* |
| External model | 0.03 | 0.03 | 0.83 | 0.408 | 0.429 |
| **BP Diastolic** |  |  |  |  |  |
| Multimodal | 0.05 | 0.02 | 2.35 | 0.019 | 0.039* |
| Gray matter | 0.02 | 0.02 | 1.00 | 0.317 | 0.354 |
| White matter | 0.04 | 0.02 | 1.73 | 0.084 | 0.113 |
| F. connectivity | 0.03 | 0.01 | 2.39 | 0.017 | 0.039* |
| External model | 0.03 | 0.03 | 1.00 | 0.318 | 0.354 |
| **Alcohol intake** |  |  |  |  |  |
| Multimodal | 0.07 | 0.02 | 2.94 | 0.003 | 0.017* |
| Gray matter | 0.08 | 0.02 | 3.73 | < 0.001 | 0.004* |
| White matter | 0.04 | 0.02 | 1.75 | 0.081 | 0.113 |
| F. connectivity | 0.03 | 0.01 | 2.02 | 0.044 | 0.077 |
| External model | 0.12 | 0.03 | 3.59 | < 0.001 | 0.077 |
| **Stroke risk** |  |  |  |  |  |
| Multimodal | 0.09 | 0.03 | 3.47 | 0.001 | 0.004* |
| Gray matter | 0.06 | 0.03 | 2.34 | 0.020 | 0.039* |
| White matter | 0.07 | 0.03 | 2.52 | 0.012 | 0.039* |
| F. connectivity | 0.03 | 0.01 | 2.38 | 0.018 | 0.039* |
| External model | 0.06 | 0.04 | 1.49 | 0.136 | 0.169 |

**Figure 5:**
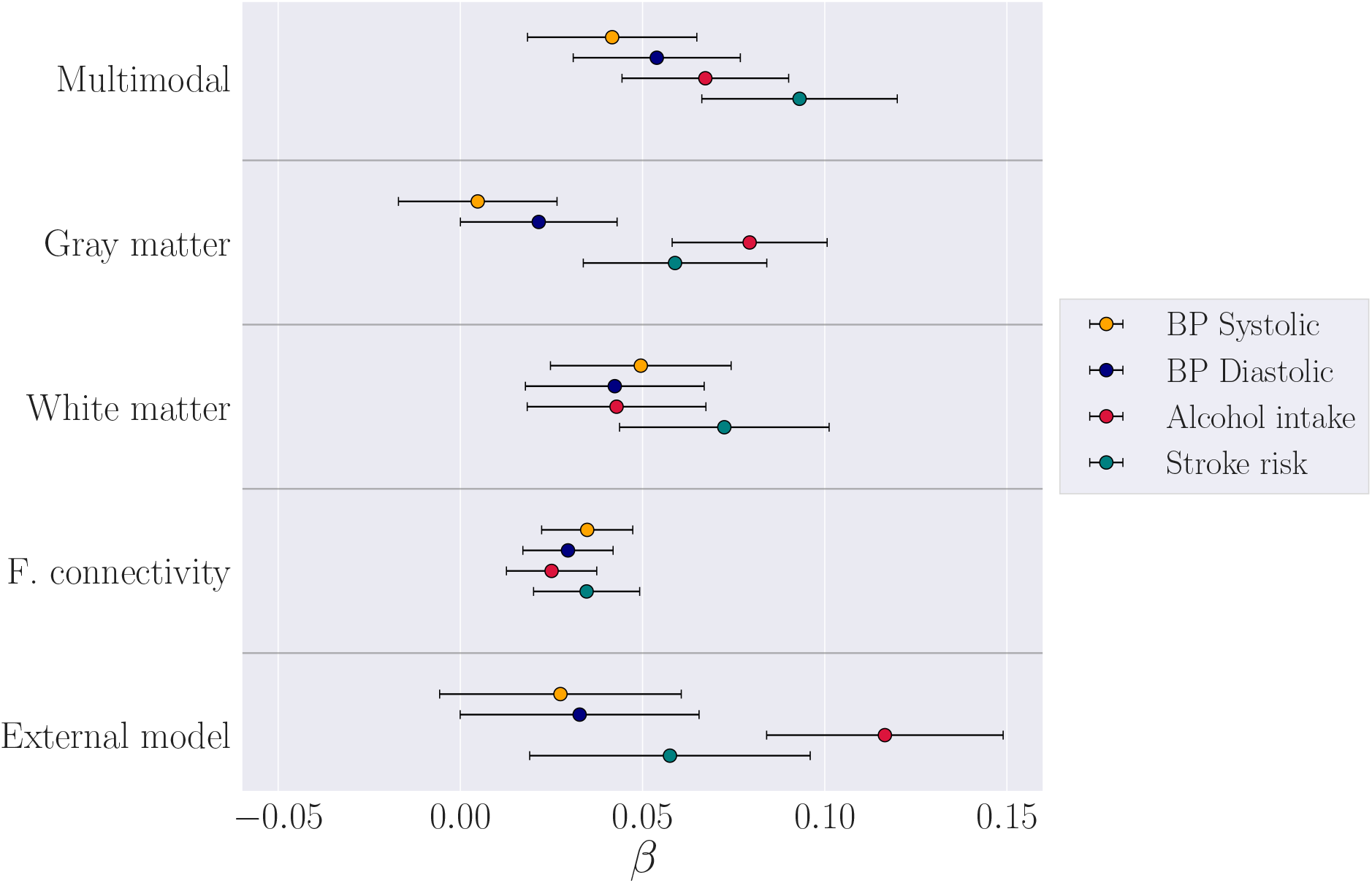
The associations (*β*± standard error) between standardized measures of brain-age delta and blood pressure, alcohol intake, and Framingham stroke risk score for each of the brain-age models. The analyses included age as a covariate.

## 4. Discussion

Our findings demonstrate that machine learning models provide meaningful imaging-based biomarkers for brain aging in healthy population cohorts [13, 15]. We found that a) the multimodal, gray matter, and white matter based models showed comparable performance, while the functional connectivity model showed lower prediction accuracy, b) blood pressure, alcohol intake, and stroke risk scores were each associated with brain-age deltas, and c) these associations were consistent across modalities.

Although previous studies have suggested better prediction with multiple imaging modalities [13, 14, 30, 54], the current study showed equivalent prediction accuracy between the multimodal model and the gray and white matter models. The exclusion of low-quality data improved the performance of the multimodal model, suggesting that established procedures for data quality control may have implications for model performance [39, 55]. Although there was also a tendency for the gray matter model to improve with the exclusion of low-quality data, discarding such data did not affect the performance of the white matter and functional connectivity models. The external gray matter model that was trained on an independent sample showed less accurate prediction of brain age in our data compared to the k-folding based gray matter model. While both datasets were corrected for factors including scanner site, motion, and ICV, such discrepancies indicate that confounding factors including recruitment procedures, scanner equipment and data-processing pipelines may influence prediction accuracy across datasets [56].

The lowest prediction accuracy was observed for rs-fMRI, indicating that the included measures of functional connectivity were less closely related to chronological age compared to structural measures. Although it is possible that voxel-wise functional connectivity measures could improve the model performance [57], the lower age-sensitivity of rs-fMRI measurements may be explained by these metrics reflecting a state, rather than trait based, assessment [58, 59, 60]. While resting-state networks, including the default mode network, are characterized as highly replicable both within-participants and across studies [45, 61, 62], investigations employing dynamic rs-fMRI conversely suggest that the connectivity between these networks may vary even within a single session of scanning [63]. Alternatively, resting-state networks and their connectivity to other networks may be preserved though plasticity despite age-related structural changes [64]. Another potential explanation is that age-related changes to network connectivity occur in a modular, rather than gradual manner [65, 66]. While gray and white matter based brain-age models provide relatively accurate predictions across studies [13, 15, 17, 19, 26], a recent application of brain-age estimation in the UK Biobank cohort similarly highlights fMRI-based brain-age prediction as a weaker correlate of chronological age (*r* = 0.434), relative to gray matter (*r* = 0.685) and DTI-based (*r* = 0.668) predictions [14].

Despite poorer performance accuracy of the functional connectivity model, the fMRI-based brain-age deltas showed associations with the biomedical variables that were similar to the other modalities. In line with recent findings from UK Biobank [14, 15], positive associations were found between brain-age deltas and diastolic blood pressure, alcohol intake, and stroke risk, concurring with previous WHII studies [32, 50], and demonstrating that the brain age-delta measure reflects individual variation in neural aging processes [30]. The associations with biomedical variables were consistent across models (see Figure 5), indicating that while modality-specific brain age models may be informative in patient groups where tissue types are differently affected by disease [26, 27, 54, 67, 68], such models may be more closely related in healthy cohorts [14]. It is possible that regional modelling of modality-specific brain aging patterns may be more suitable to detect specific associations with biomedical and clinical measures [19], which could get lost in machine learning models that summarise aging across the whole brain to produce a single global prediction [27].

In conclusion, machine-learning based brain age prediction can reduce the dimensionality of neuroimaging data to provide meaningful biomarkers of individual brain aging. While the presented imaging-derived markers can help to assess general effects of clinical and biomedical risk factors on the brain, models of distinct and regional neural aging patterns may result in more refined biomarkers that can capture additional biological detail [19, 26, 27].

## Acknowledgements

We thank all Whitehall II participants for their time. Work on the Whitehall II MRI sub-study was funded by the “Lifelong Health and Wellbeing” Programme Grant: *“Predicting MRI abnormalities with longitudinal data of the Whitehall II Substudy”* (UK Medical Research Council: G1001354, PI: K.P.E), and the HDH Wills 1965 Charitable Trust (Nr: 1117747, PI: K.P.E). This research received funding from the the NIHR Oxford Health Biomedical Research Centre (N.F), the Research Council of Norway (AMG.dL, T.K., L.T.W.; 286838, 273345, 249795, 276082, 223273), the South-East Norway Regional Health Authority (L.T.W.; 2015073, 2019107), the European Research Council (ERC) under the European Union’s Horizon 2020 research and innovation programme (L.T.W.; Grant agreement No. 802998) and a UKRI Innovation Fellowship (J.H.C.; MR/R024790/1). The use of data from the UK Biobank was approved by the UK Biobank Access Committee (Project No. 27412).

